# Modelling avoidance in pathologically anxious humans using reinforcement-learning

**DOI:** 10.1101/081984

**Authors:** Anahit Mkrtchian, Jessica Aylward, Peter Dayan, Jonathan P Roiser, Oliver J Robinson

## Abstract

Serious and debilitating symptoms of anxiety are the most common mental health problem worldwide, accounting for around 5% of all adult ‘years lived with disability’ in the developed world. Avoidance behaviour –avoiding social situations for fear of embarrassment, for instance–is a core feature of such anxiety. However, as for many other psychiatric symptoms, the biological mechanisms underlying avoidance remain unclear. Reinforcement-learning models provide formal and testable characterizations of the mechanisms of decision-making; here, we examine avoidance in these terms. One hundred and one healthy and pathologically anxious individuals completed an approach-avoidance go/no-go task under stress induced by threat of unpredictable shock. We show an increased reliance in the anxious group on a parameter of our reinforcement-learning model that characterizes a prepotent (Pavlovian) bias to withhold responding in the face of negative outcomes. This was particularly the case when the anxious individuals were under stress. This formal description of avoidance within the reinforcement-learning framework provides a new means of linking clinical symptoms with biophysically plausible models of neural circuitry and, as such, takes us closer to a mechanistic understanding of pathological anxiety.

## Introduction

Avoidance is a core feature of anxiety^1^ and plays a central role in psychological strategies for the treatment of anxiety^2^, but its underlying neural and cognitive mechanisms are unknown. Avoidance can be adaptive: if an individual perceives a situation as stressful then it makes sense to avoid that stressor in the future. However, excessive avoidance can result in a pathological downward-spiral. The more one avoids a situation, the less opportunity there is to learn that the situation is not as bad as feared, and a vicious cycle of avoidance and anxiety emerges^1^. For example, an individual who fears social embarrassment might ultimately end up housebound, avoiding all social interaction.

The diathesis-stress model of anxiety^3^ proposes that maladaptive avoidance should be greatest during periods of environmental stress in vulnerable individuals. This idea has clear face-validity, and is supported by clinical anecdote, but is largely derived from retrospective, subjective self-report. This is because quantifying avoidance under stress in an experimentally controlled yet ecologically valid manner in humans is methodologically challenging. In this study we address this challenge using: i) a translationally-validated ‘threat of shock’ procedure to induce stress^4^,^5^; ii) a cognitive task that has been shown to reliably index avoidance behaviour in healthy individuals^1^; and iii) a computationally precise method of defining of avoidance.

Specifically, we operationalize avoidance as a behavioural bias towards withholding action (“no-go”, i.e. inhibition) in the face of potentially negative outcomes. This powerful prepotent reflexive (or Pavlovian) bias has been observed consistently in humans and animals ^6^–^9^ and is so profound that it can disrupt instrumental goal-directed behaviour ^6^–^9^. This is known as Pavlovian-Instrumental transfer^10^, and we harness it here to measure the degree to which individuals rely on their prepotent avoidance biases.

Reinforcement-learning algorithms can provide parameterizations of avoidance behaviour that offer insight into both optimal behaviour when set correctly^11^, and to dysfunction and pathology when set incorrectly^12^. Critically, reinforcement-learning models enable us to parameterize the influence of Pavlovian avoidance biases on task performance in a formal manner. A large body of work has applied these models to healthy humans ^6^–^8^, and they form the basis of human-level artificial intelligence^11^, but to date they have not been applied to individuals with anxiety disorders.

We therefore tested pathologically anxious and healthy individuals completing an approach-avoidance go/no-go task under stress, which was induced by threat of shock. Avoidance was defined and parameterised within a reinforcement-learning framework. We predicted that anxious individuals would show high reliance on avoidance bias, and that this would be exacerbated by stress.

## Materials and Methods

### Participants

All data, task scripts and code to recreate the figures in this paper are freely available online^*^. A total of 101 participants were included in the study. Healthy participants (N=58 (originally N=62 but four excluded because they failed to follow task instructions); 36 male [62.1%]; age range: 18-57; mean (standard deviation) age=26.7 (7.1)) and (minimum 6 month) unmedicted individuals suffering from pathological anxiety symptoms (N=43; 27 male [62.8%]; age range 18-53; mean age=28.8 (8.8)) were recruited from online advertising and institutional subject databases. The primary difference between the groups in initial recruitment was that only the pathological group self-defined as experiencing distress from mood/anxiety symptoms. Healthy participants responded to an advertisement asking for healthy individuals with no psychiatric symptoms. A phone screen confirmed no history of psychiatric, neurological or substance use disorders. The pathological group responded to an advertisement for individuals suffering with low mood, anxious or depressive symptoms. Following an initial phone screen, individuals who met criteria for mood or anxiety disorder symptomatology according to a face-to-face Mini International Neuropsychiatric Interview (MINI)^13^ were included. According to the MINI, the majority of participants (N=27) met criteria for both GAD and MDD (N=9 with additional panic disorder), N=8 met criteria for GAD (N=3 with panic disorder, N=1 with agoraphobia), N=2 Panic disorder and MDD, and a further N=6 MDD alone. The average number of depressive episodes was 5 (standard deviation ±7).

Exclusion criteria were any form of medication within the last 6 months, any current psychiatric diagnosis (other than major depression or anxiety disorder), neurological or substance use disorders, or pacemaker. Previous unsuccessful treatments greater than 6 months prior to testing included medication (N=5), psychological treatment (N=8) or both medication and psychological treatment (N=12). A small number were undergoing current psychological treatment (N=5) but still met diagnostic criteria and 1 participant with MDD narrowly missed the diagnostic threshold (according to the MINI) but was included due to clear evidence of distress. A measure of trait anxiety was obtained for all participants using the State-Trait Anxiety Inventory (STAI)^14^. All participants provided written informed consent and were reimbursed £7.50/hour for participation. The study obtained ethical approval from the UCL Research Ethics Committee (Project ID Numbers: 1764/001 and 6198/001).

Sample size was determined using *a priori* power calculations. The healthy control sample was powered for an effect size of d=0.49 based on the t-test of the within-subjects effect of an anxiogenic manipulation on a different action valence task^15^: N=57 gives 95% power for a two-tailed t-test with α = 0.05. The size of the pathological group was based on an assumed between-groups effect size of 1.09 (observed in our prior study: ^16^), which was decreased to 0.8 for the purpose of a conservative power analysis. N=42, gives 95% power for a two-tailed between-groups t-test with α = 0.05. Non computational analyses were completed using JASP.^17^

### Manipulation

State anxiety was induced via threat of unpredictable electric shocks delivered with two electrodes attached to the non-dominant wrist using a Digitimer DS5 Constant Current Stimulator (Digitimer Ltd, Welwyn Garden City, UK). A highly unpleasant (but not painful) subjective shock level was established using a shock work-up procedure prior to testing. No more than five (to avoid habituation) shocks with gradual increasing shock level were administered. Participants rated each shock on a scale from 1 (barely felt) to 5 (unbearable). Shock level was matched at a level of 4 across participants. The experimental task was programmed in Psychtoolbox (http://psychtoolbox.org) for MATLAB 2014 (The MathWorks Inc., Natick, MA), presented on a laptop and administered under alternating safe and threat blocks. During the safe block, the background colour was blue and proceeded by a 4000ms message stating: “YOU ARE NOW SAFE FROM SHOCK”. During the threat block, the background colour was red and the message: “WARNING! YOU ARE NOW AT RISK OF SHOCK” was presented for 4000ms. Participants were told that they might receive a shock only during the threat condition but that the shocks were not dependent on their performance. In practice, a single shock was delivered at a pseudorandom timepoint during one-third of threat blocks (a total of four shocks across 480 trials). Note that it is the anticipation of these shocks, not the shocks themselves that constitutes the manipulation (see supplemental analysis). At the end of each experimental task, participants retrospectively rated how anxious they felt during the safe and threat conditions on a scale from 1 (“not at all”) to 10 (“very much so”).

### Approach-Avoidance Task

The task was based on the design of a previous probabilistic go/no-go reinforcement learning task^8^,^15^ modified to incorporate the threat manipulation. The task comprised four experimental conditions where action (go/no-go) was crossed with valence (reward/punishment): 1) go to win reward (GW), 2) go to avoid losing (GA), 3) no-go to win reward (NGW), and 4) no-go to avoid losing (NGA). On each trial, participants were presented with one of four fractal cues per condition, followed by a target detection task, and subsequently by a probabilistic outcome (Figure 1). The fractal cue, target detection task and the outcome were each presented for 1000ms and separated by a 250ms inter-trial interval (ITI). Each fractal cue signified one of the four experimental conditions, but this was not made explicit at the start of the experiment. Thus, subjects had to learn that each fractal image indicated both which 1) action (go=make response; no-go=withhold response) to perform during the target detection task and 2) the associated valence of the outcome (reward/no reward; punishment/no punishment). The meaning of the fractal cues was randomised across participants. In the target detection task, a circle was presented randomly on one side of the screen (50% of trials on the left). In the go experimental conditions (GW/GA), participants had to match the position of the circle by pressing the corresponding key (i.e., press the left key when the circle was on the left and vice versa). In the no-go experimental conditions (NGW/NGA), participants had to withhold any response (i.e. any response was recorded as incorrect). The circle was presented for 1000ms regardless of response.

In the rewarded conditions (GW/NGW), correct responses were rewarded 80% of the time, but resulted in no win 20% of the time. Incorrect responses led to no win 80% of the time, but were rewarded 20% of the time. In the punishment conditions (GA/NGA), correct responses avoided punishment 80% of the time but led to a loss 20% of the time (and vice versa for incorrect responses). Wins were indicated by a happy face and a gain of 10 points. Losses were indicated by a fearful face and a 10 point deduction. A horizontal yellow bar indicated when participants neither won nor lost points. Faces were selected from the Ekman facial set and the genders of the faces were counterbalanced across participants.

Participants were informed about the probabilistic nature of the task but they were not told the action-outcome contingencies for each fractal cue. Instead, they were told that they had to learn the correct response for each fractal cue, which could be either a go response or a no-go response, by trial and error. The task was divided into 24 alternating safe and threat blocks (12 blocks of each) with the order of the safe and threat conditions counterbalanced across participants. A different set of fractal cues was used under threat and safe in order to avoid possible confounding effects from learning under the different conditions. The eight fractal cues for threat and safe (four in each condition) were counterbalanced across participants.

Each block had five trials per experimental condition (GW, GA, NGW, NGA), with a total of 20 trials per block. The trials were randomly presented within each block. There were thus a total of 240 trials (60 trials for each fractal cue) per safe or threat condition. The task lasted around 35 min with a single shock delivered in the third, seventh, tenth and twelfth threat blocks. These shocks were always presented in the ITI between trials (the 4^th^ trial of the 3^rd^ threat block, the 18^th^ trial of the 5^th^ threat block, the 10^th^ trial of the 10^th^ threat block and the 2^nd^ trial of the 12^th^ threat block). Critically, these shocks were presented to maximise manipulation efficacy^18^ (see supplemental analysis of effect on choice behaviour). Prior to the start of the task, participants completed nine practice trials without the threat manipulation. Each outcome appeared three times and identical black images were used instead of fractal cues in order to familiarise participants with the task without confounding learning of the action-outcome contingencies.

### Reinforcement-learning models

Reinforcement-learning modelling proceeded in the same way as described in a prior paper^8^. Briefly, we built seven parameterized reinforcement-learning models to fit to the behaviour of the subjects. All models were adapted Rescorla Wagner models. We use the term *‘Standard’* to denote the 6 parameter winning model from Guitart-Masip, et al. (2012) and either add or subtract parameters to test model fits for seven separate models (See Table 1 for a parameter specification summary).

**Table 1:** Model specification (NP = number of parameters)

| Model Name | NP | Parameters |  |  |  |  |  |  |  |
| --- | --- | --- | --- | --- | --- | --- | --- | --- | --- |
| *Standard - Action Bias* | 5 | Reward sensitivity | Punishment sensitivity | Learning rate |  | Lapse | - | Approach-Avoid bias |  |
| *Standard – Approach-Avoid* | 5 | Reward sensitivity | Punishment sensitivity | Learning rate |  | Lapse | General action bias | - |  |
| *Standard + 2 Approach-Avoid - 1 Sense* | 6 | Sensitivity |  | Learning rate |  | Lapse | General action bias | Approach bias | Avoidance bias |
| *Standard* | 6 | Reward sensitivity | Punishment sensitivity | Learning rate |  | Lapse | General action bias | Approach-Avoid bias |  |
| *Standard + 2 Approach-Avoid* | 7 | Reward sensitivity | Punishment sensitivity | Learning rate |  | Lapse | General action bias | Approach bias | Avoidance bias |
| *Standard + 2 Learning Rates* | 7 | Reward sensitivity | Punishment sensitivity | Reward Learning rate | Punishment Learning rate | Lapse | General action bias | Approach-Avoid bias |  |
| *Standard + 2 Approach-Avoid + 2 Learning Rates* | 8 | Reward sensitivity | Punishment sensitivity | Reward Learning rate | Punishment learning rate | Lapse | General action bias | Approach bias | Avoidance bias |

### Learning models

All the models assigned a probability to each action a_t_ on trial t based on an action weight and the current stimulus. The action weights were constructed according to a simple Rescorla–Wagner-like update equation with a learning rate. Reinforcements were coded as +1 for a reward, -1 for a punishment and 0 for no feedback. A sensitivity parameter determined the effective size of reinforcements for a subject. For the majority of models the sensitivity parameter could take on different values for the reward and punishment trials. For one model (*‘Standard + 2 Approach-Avoid - 1 Sense’*) there was only one sensitivity parameter per subject, thus assuming that failure to obtain a reward was as aversive as obtaining a punishment. The initial value for the go action was set to zero and the action weight was modified to include a static general action bias parameter which denoted overall go tendency (with the exception of one model *‘Standard - action bias’*, in which this was not included). The Pavlovian approach-avoid bias parameter (excluded for one model *‘Standard - Approach-Avoid*) inhibited the tendency to go in proportion to the negative value of the punishment stimulus, while it similarly promoted the tendency to go in proportion to the positive value of the reward stimulus. For the model with two *Approach-Avoid* parameters (*‘Standard + 2 Approach-Avoid*), there were two parameters, updated separately for rewarded and punished trials. For the models with two learning rates (‘*Standard + 2 Approach-Avoid + 2 learning rates’* / ‘*Standard + 2 learning rates’*), there were separate learning rates for rewarded and punished trials. In sum, for a given action (a=go / no-go), stimulus (s=GW / GA / NGW / NGA), reinforcement (r=+1 /-1 / 0) on each trial (t):

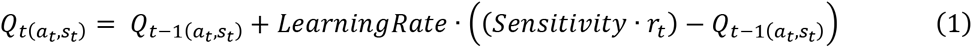

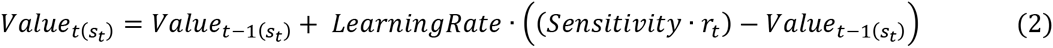

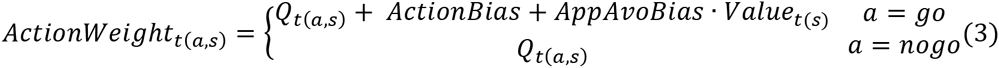

### Observation model

For action selection, the probability of each action was passed through a squashed softmax function with the addition of an irreducible lapse parameter (referred to as ‘noise’ in earlier papers, but renamed lapse here to avoid confusion with temperature noise parameters), which was free to vary between 0 and 1.

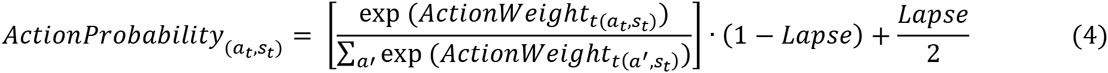

## Parameter Estimation

We used an hierarchical Type II ML expectation–maximization (EM) procedure to fit the parameters across all subjects and conditions. These procedures are identical to those used by Huys et al 2011^10^. Briefly, the top level of the hierarchical model specified distributions over the parameters for the subjects (see below). At each iteration, the current top-level distributions were used as a prior for a Laplace approximation to the intermediate-level posterior distribution of the parameters for each subject (the E-phase). These intermediate-level distributions were then used to determine the next iteration of the top-level distributions (the M-phase). The algorithm was initialized with maximum likelihood values of all the parameters for the subjects; the Laplace approximation was based on the use of fminunc in MATLAB, using multiple random initial values at each iteration of optimization to help avoid local minima. Four different population distributions were tested (see Figure 3):

1. Four distributions: one for anxious individuals under threat, one for controls under threat, one for anxious individuals under safe, one for controls under safe. This is the most relaxed procedure and serves to pull all parameters apart.
2. Two distributions: one distribution for threat and one distribution for safe. This fitting procedure was blind to the existence of group.
3. A single distribution for all participants and conditions (i.e. each participant was included twice within the distribution; once for the safe, and once for threat conditions). This fitting procedure was blind to the existence of both group and threat condition, and serves to pull all parameters closer together.
4. Two distributions: one distribution for anxious individuals and one distribution for controls. This fitting procedure was blind to the existence of induced anxiety.

The fit of each model and distribution was compared using the integrated BIC (iBIC). The iBIC is the integral of the likelihood function over the individual parameters (for details, see^10^). Small iBIC values indicate a model that fits the data better *after* penalizing for the number of parameters. The parameter fitting procedure results in one iBIC per distribution. These are then summed together to provide a single iBIC to enable model comparison across distributions. The lowest overall iBIC denotes the ‘winning’ model and distribution combination (an approximate Bayes Factor of the comparison of iBIC scores can be calculated using exp(ΔiBIC/2).) Note that fitting the parameters of the winning model using a different, hierarchical Bayesian, approach recovered similar parameters (see supplement).

The parameters recovered from the winning model were then compared across groups and conditions using permutation tests implemented in MATLAB 2014 (see http://tiny.cc/2slwby for code and supplement for replication using different software). The recovered p-values are comparable to those derived from standard t-tests, but do not require the assumption of normality (critical given the possibility of multimodal distributions recovered from the model fitting procedure).

## Results

### Basic analysis of symptoms and behaviour

Our sample consisted of 58 healthy individuals and 43 unmedicated individuals suffering from pathological anxiety symptoms. The learning task (adapted from^8^,^15^) comprised four experimental trial types which orthogonalized action (go/no-go) and valence (reward/punishment; see Figure 1 for details). All trials were completed under threat of shock (i.e. stress) and safe conditions.

**Figure 1.**
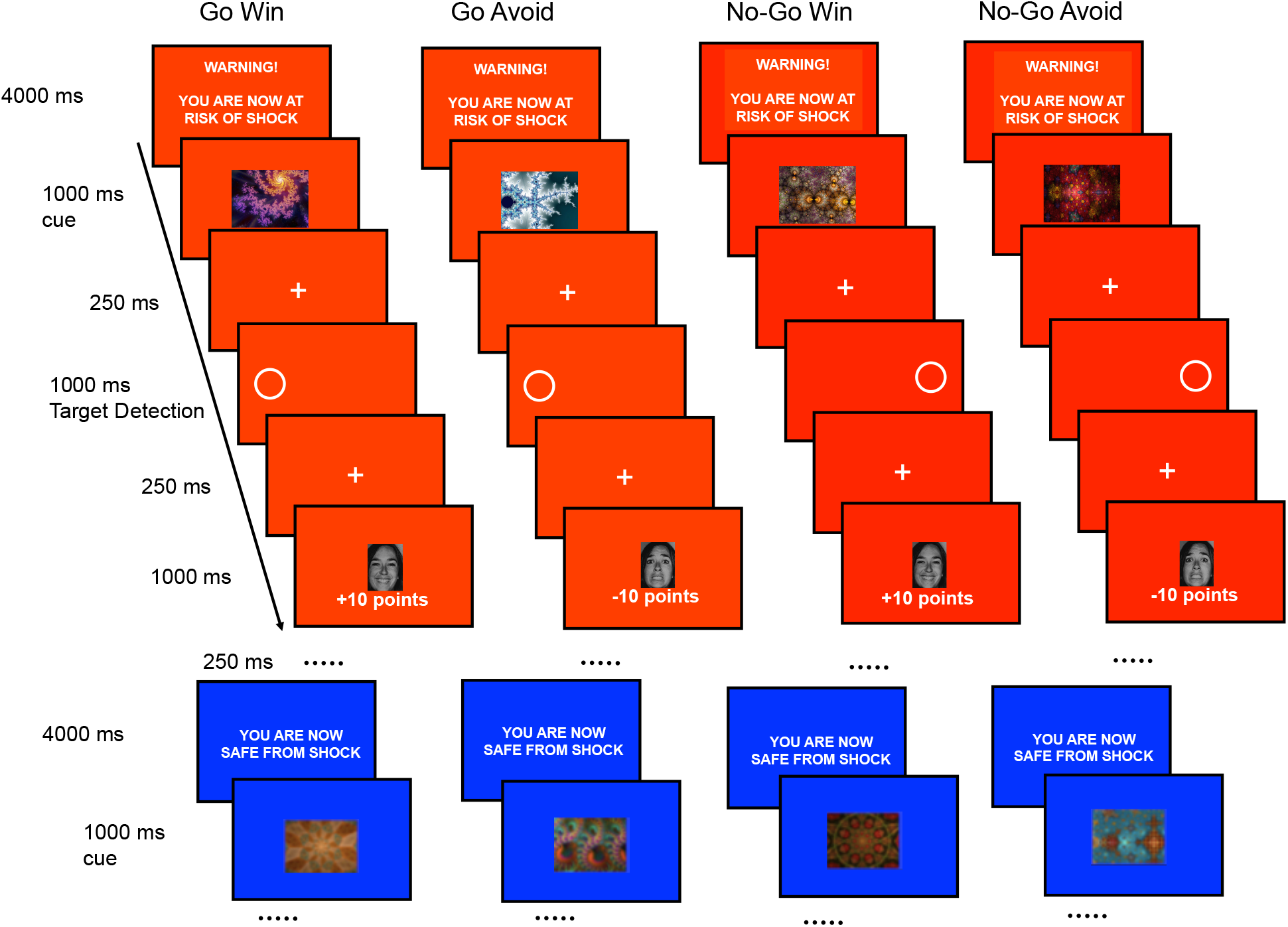
Experimental paradigm: *The trial sequence for each trial-type condition under threat (red) and safe (blue) conditions. There were equal numbers of Go to Win (GW), Go to Avoid (GA), No-Go to Win (NGW) and No-Go to Avoid (NGA) trials within each safe and threat block, and these were randomly ordered within each block (note that safe sequence proceeds in the same way as threat but is curtailed here for brevity). The prepotent Pavolovian bias to a win is a go response (approach) and the prepotent Pavlovian response to a loss is no-go (avoid); hence in GW and NGA, the bias and task instructions are aligned; but in G A and NGW, participants have to learn to overcome their avoidance and approach biases respectively. The safe and threat blocks were presented in alternating order, counterbalanced across participants. A different set of fractal cues was used for the safe and threat blocks, counterbalanced across participants. At feedback, a face (happy +10 points, fear -10 points) was shown 80% of the time, and no points (i.e. a yellow bar – not shown in the figure) was shown 20% of the time.*

As expected, the anxious group reported significantly higher symptoms of trait anxiety relative to controls (F(1,97)=69.9, η_p_^2^=0.4, p<0.001); Figure 2a). Participants retrospectively reported feeling greater anxiety during the stress manipulation relative to the matched safe condition (F(1,99)=166, η_p_^2^=0.6, p<0.001; Figure 2b), which was similar between groups (main effect of group: F(1,99)=2.0, η_p_^2^=0.02, p=0.16; group x condition interaction F(1,99)=0.007, η_p_^2^<0.001, p=0.9).

Analysis of overall performance accuracy revealed a main effect of action (F(1,99)=90, η_p_^2^=0.5, p<0.001), qualified by an action (go/no-go)-by-valence (reward/punishment) interaction (F(1,99)=94, η_p_^2^=0.5, p<0.001; Figure 2c). As expected, this was driven by worse performance in the conditions where Pavlovian biases had to be overcome in order to make the appropriate response (i.e. a loss-driven avoidance bias in GA; and a win-driven approach bias in NGW) as well as an overall bias towards making go responses. There was a main effect of group (F(1,100)=15, η_p_^2^=0.1, p<0.001) driven by worse overall accuracy in anxious individuals, but no other interactions with group or condition (all p>0.5). However, as apparent in Figure 2d, learning follows a complex time-course which differs by condition (and by individual). We therefore turned to a computational model-based analysis to examine these differences at a fine scale.

**Figure 2:**
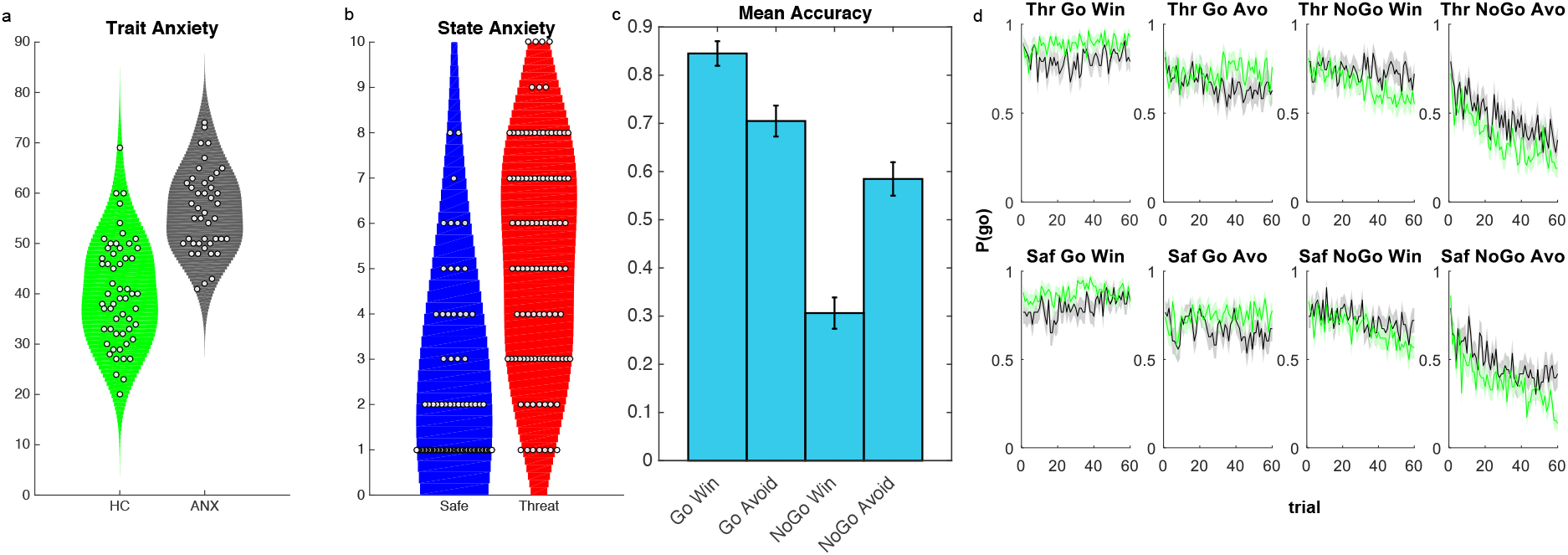
Self-report anxiety and task performance. *Between groups, a) our anxious sample reported significantly higher trait anxiety scores (data missing for two participants in the healthy group), while b) the whole sample reported increased (induced) anxiety, rated retrospectively, under threat relative to safe conditions (violin plots; each point represents a subject, background shading represents estimated distribution). c) Collapsed mean accuracy differs as a function of trial type, but this ignores that d) performance on the task changed over time such that the probability of making a response (P(go)) differed as a function of trial type, condition and group (shading represents standard error of the mean). (HC=healthy control=green; ANX=pathological anxiety=grey; Saf =safe; Thr= threat; Avo=Avoid)*

### Reinforcement-learning model selection and validation

We fitted reinforcement-learning models to trial-by-trial choice behaviour using an hierarchical Type II maximum likelihood expectation–maximization approach^10^. The most parsimonious model (*‘Standard + 2 Approach-Avoid + 2 Learning Rates’*; Table 1; Figure 3e; methods) is an adapted Rescorla-Wagner model^19^ identical to the winning model in prior studies of healthy individuals^6^,^8^, with the exception that there are separate Pavlovian approach, avoid and learning rate parameters for the cases of rewards and punishments. In other words, this model included an approach bias parameter, an avoidance bias parameter, and accommodated separate speeds of learning about rewards and punishments.

**Figure 3:**
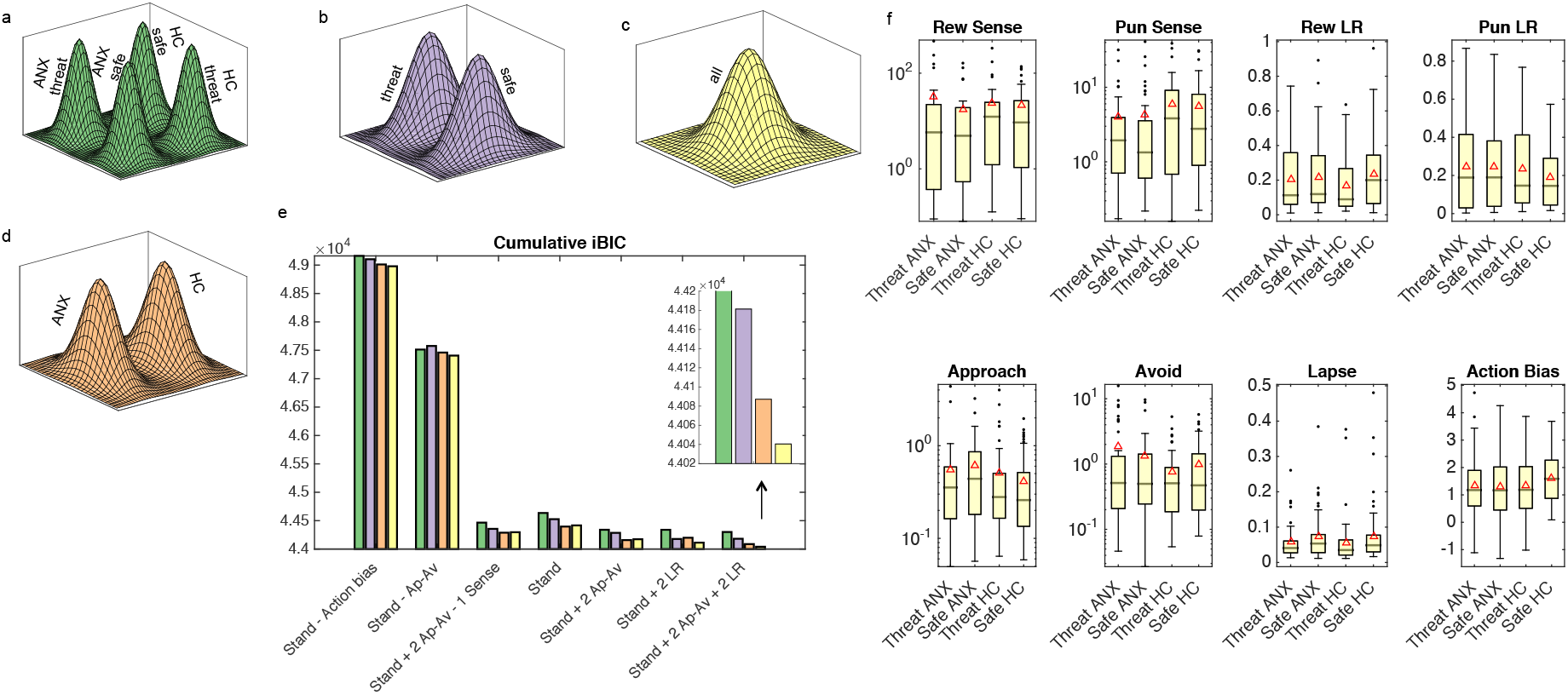
Model Fitting and Comparison. *Four different population distributions were tested separated by a) group and threat condition (4 distributions); b) by threat condition alone (2 distributions); c) blind to group and threat condition (1 distribution); and d) by group alone (2 distributions). Comparison of models and distributions using integrated Bayesian Information Criteria (iBIC) scores (colours match distributions throughout figure) revealed a winning model of ‘Standard + 2 Approach-Avoid + 2 Learning Rates’, fit across a single prior distribution (inset zoomed in on the distribution comparison for this model). Box and whisker plots of the recovered parameters from the wining model/distribution are presented in f) separated by group and condition (red triangles denote means, lines denote medians; based on individual parameter estimates). Note that transforming the data into interpretable space skews the distributions of inferred parameter values away from normality and thus many values fall outside the whiskers. Log scales are used for the sensitivity and approach-avoidance parameters to aid visualisation of these exponentially transformed parameters (HC=healthy control; ANX=pathological anxiety; Rew=reward; Pun=punishment; Stand= standard; Ap-Av=approach avoid; Sense=sensitivity; LR=learning rate; Avoid=Avoidance Bias; Approach=Approach Bias)*

The hierarchical model fitting procedure requires the specification of population level priors. This raises an important conceptual question when it comes to considering multiple groups. Should we consider groups of anxious individuals and healthy controls as being sampled from the same or different populations? We answered this question through the adoption of a population-level model comparison approach. We compared fits for models ranging from four separate prior distributions for each group and stress condition (Figure 3a) to a single distribution for all subjects and conditions (Figure 3c). The best fit for our winning model was achieved by fitting a single population distribution (Figure 3c), implying that we did not obtain sufficient evidence to suggest that anxious and healthy individuals were sampled from different populations. Box plots and means of the posterior parameter distribution across subjects (under the Type II empirical prior) are shown in Figure 3f; that all subjects share the same prior implies that the recovered parameters will be drawn closer together.

We next ran a posterior predictive model with parameters set to those from the winning model (i.e. having a computer make decisions as if it was each individual subject). Average parameters recovered from simulated data were close to those that were originally observed (Figure 4a), albeit with more noise for the NGW condition. Average simulated behaviour over time matched closely that of the subjects (Figure 4b; compare to Figure 2c)

**Figure 4:**
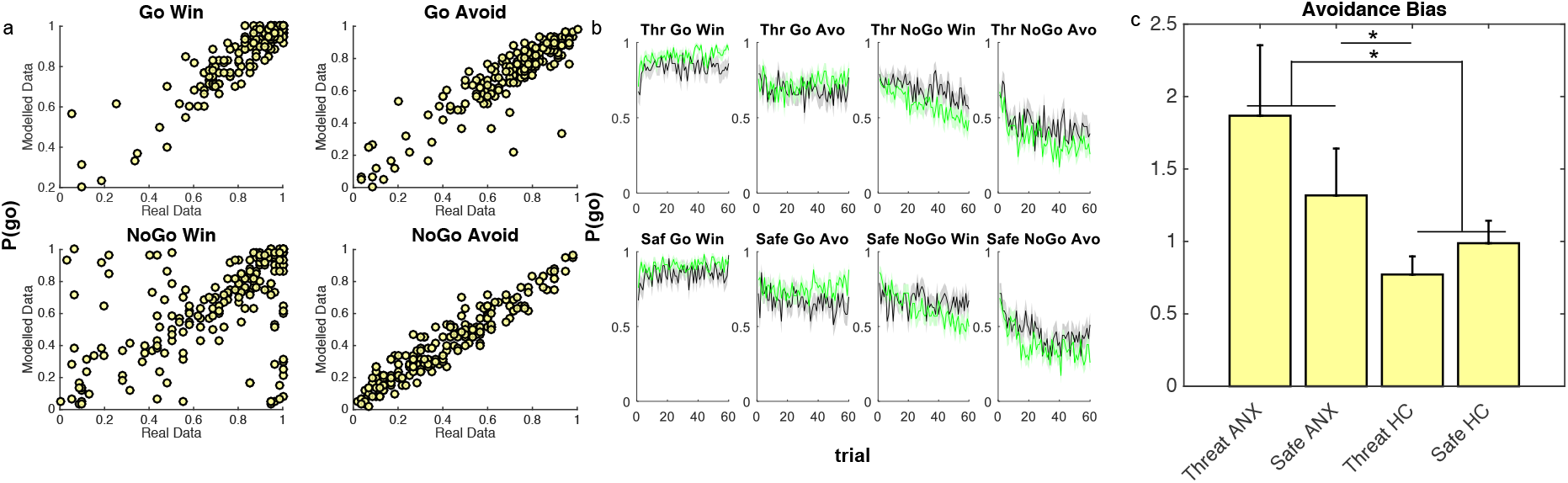
Posterior Predictive Model. *Running the estimated parameters for each subject through a posterior predictive model recovered both a) average go probabilities for each trial type (sensitivity plots: each marker represents one subject under one condition so there are twice as many markers as subjects), and b) group-averaged trial-by-trial performance (compare to real data in Figure 2c). Comparing parameters across group and condition revealed c) a significantly higher avoidance bias parameter in pathological anxiety across conditions as well as greater threat-potentiated avoidance in pathological anxiety (error bars represent standard error of the mean; HC=healthy control=green; ANX=pathological anxiety=grey; Saf=safe; Thr=threat; Avo=Avoid)*

### Pathological anxiety is associated with increased reliance on avoidance bias, especially under stress

We finally performed permutation tests on the posterior parameters to assess the effects of group and threat condition. These revealed an increased reliance on the avoidance bias parameter in the anxious group (effect of group averaged across threat and safe): P_(permutation)_=0.03) (Figure 4c) and a significantly greater increase in the avoidance parameter under threat vs safe conditions in the anxious group relative to controls (P_(permutation)_=0.012; Figure 4d) driven by a significantly greater avoidance in anxious relative to controls under threat (P_(permutation)_=0.008), but not safe (P_(permutation)_=0.18) conditions. No other parameter differed across either group or condition (all P_(permutation)_>0.1).

## Discussion

Anxious individuals show strong avoidance behaviour that can be debilitating and self-perpetuating^1^. Here, using a novel computational approach, we provide evidence that pathologically anxious individuals show increased reliance on an avoidance bias (a Pavlovian bias to withhold responding in the face of punishments) during reinforcement-learning. Moreover, consistent with the diathesis-stress hypothesis, this effect was exacerbated under stressful conditions only in the anxious group.

We provide a potential computational mechanism for this effect. We show that avoidance behaviour – which is currently measured by retrospective self-report - can emerge at the level of stimulus-action associations. Specifically, anxious individuals may show avoidance in the face of threats because they inhibit their action tendencies when faced with a perceived negative outcome. Over time, individuals may be ultimately able to learn to overcome this bias (i.e. promote instrumental override of Pavlovian bias parameters) if they are given the opportunity to experience outcomes (i.e., NGW go probability is lower at the end than GW here). However, in the real world, avoidance means that, by definition, predicted outcomes are rarely experienced and challenged, there is little opportunity to learn, and a persistent miscalibration can emerge.

The growing field of computational psychiatry^12^ seeks to use theory-driven approaches to explain psychiatric phenomena. Testable theories are a pre-requisite to a clear mechanistic understanding: here, we have outlined a precise and formalised computational theory about how avoidance emerges in anxiety under stress. This approach has at least two further advantages. Firstly, it allows us to reduce a highly dimensional dataset (here, choices over time) into small number of parameters that respect the temporal variability of the data (unlike responses averaged over time). Secondly, we can directly integrate this model into biophysically plausible models of underlying neural activity^20^. Indeed, performance of this task in healthy individuals has been linked neurocognitively to striatal and midbrain regions associated with network models of action^7^,^8^ as well as dopaminergic modulation of this circuitry^21^. Striatal regions of this circuitry are also modulated by the threat of shock technique used here^18^, providing a link between these substrates and stress. This computational approach therefore holds promise as a means of unifying complex psychiatric phenomena, such as avoidance, with their underlying neural circuitry.

Such a mechanistic link is critical if we wish to develop improved treatments. Without mechanistic understanding, treatment development has to be targeted at downstream symptoms – e.g. self-reported avoidance. The problem with this approach can be illustrated by the symptom of cough^22^. Lung cancer, allergies, bronchitis or tuberculosis all result in a cough through fundamentally different mechanisms, but the treatment for one will be ineffective for the others (and indeed may even cause harm through side effects). Targeting clearly defined mechanisms, not symptoms, should ultimately improve the effectiveness of interventions. For example, extensive work in the development of psychological interventions for anxiety has suggested that exposure therapy should be paired with behavioural training to overcome avoidance in order to be effective^1^, but the mechanism is unclear. The present findings suggest that this may be because such training encourages an instrumental override of Pavlovian bias during action selection. One avenue for future exploration, therefore, is whether training to overcome bias on GA trials on tasks like the present could promote instrumental override (cf.^23^ but also^24^). If proven effective, such speculative task-based interventions (completed via smartphones, for example) could have enormous potential value for public health.

## Acknowledgements

This research was funded by a Medical Research Council Career Development Award to OJR (MR/K024280/1) and a Medical Research Foundation Equipment Competition Grant (C0497, Principal Investigator OJR). We thank Quentin Huys and Woo-Young Ahn for parameter fitting code and assistance.

## Contributions

OJR conceived the experiment. OJR and AM designed the study with critical input from JPR. AM programmed the task. AM and JA screened participants, collected the data and performed analyses under the supervision of OJR. OJR performed the computational modelling under the supervision of PD. OJR wrote the paper with critical input from JPR, PD, JA and AM.

## Supplement

### Effect of shocks

Comparing pooled performance on the five trials before and after the four shocks (i.e., 20 pre-and 20 post-shock trials) revealed no impact of the shock stimulation on accuracy (pre- vs post-shock: F(1,99)=1.6, p=0.21, η_p_^2^=0.016; pre- vs post-shock*group: F(1,99)=0.7, p=0.39, η_p_^2^=0.007) or reaction time (pre- vs post-shock: F(1,98)=0.13, p=0.72, η_p_^2^=0.001; pre- vs post-shock*group: F(1,98)=0.1, p=0.75, η_p_^2^=0.001). An analysis separate by trial type (GA, NGA, GW, NGW) revealed no significant interactions (all p>0.2).

### Model-fitting

Fitting our winning model using a hierarchical Bayesian approach implemented using the hBayesDM (hierarchical Bayesian modeling of Decision-Making tasks) toolbox^1^ recovered very similar avoidance parameters (correlation between parameters: avoidance under threat: r=0.85, p<0.001; avoidance under safe: r=0.84, p<0.001).

We note that our model does a better job of fitting the trials that contribute to the avoidance bias parameter fitting (i.e. the avoid trials; Figure 4a) than the rewarded trials (especially NGW). This means that inference is based on the trials that are best captured by the model. Future work might seek to refine model components that improve the fit of the rewarded trials. One option is to test the effect of varying the population priors at the parameter level, rather than the model level. For example it may be that avoidance bias is best fit across a single population, but that other parameters would be best fit across separate populations. Specifying such differences is out of reach of our current tools, but should become available within the coming years. Another potential way to improve fits would be to run permutation tests at the model-fitting stage (rather than on the final recovered parameters). In other words, trial-by-trial data could be randomly assigned to groups and conditions prior to fitting, with as many permutations as possible tested. With our current computing setup this procedure would take several hundred hours to run, but this approach may become feasible within the coming years.

### Group parameter comparisons check

We confirmed our permutation tests with independent Matlab code (http://tiny.cc/oygmdy) and the R package coin (http://tiny.cc/o6brdy *IndependenceTest*). Both showed increased reliance on the avoidance bias parameter in the anxious group (effect of group on average across threat and safe: p(permutation_matlab)=0.044 / p(permutation_R)=0.049 and a threat*group interaction: (p(permutation_matlab)=0.015/ p(permutation_R)=0.018.

## Footnotes

* * https://figshare.com/articles/Avoidance_Anxiety_Materials/3860250

